# Deciphering the limitations of immortalized hepatocyte cell lines for the study of liver cis-regulatory elements

**DOI:** 10.64898/2026.06.05.730479

**Authors:** Andrew Bellesis, Xinyi Li, Dervla Moore-Frederick, Deyuan Xu, Kara Delbridge, Junjie Ma, Gabriella Vaccaro, Buddhima Athukorala Aracchige Edward, Maddie Kellogg, Yehuda Creeger, Alexander S. Okamoto, Irene M. Kaplow

## Abstract

**Background:** Immortalized cell lines are widely used to study transcriptional regulation. However, previous studies have shown that immortalization and prolonged culture can alter their regulatory landscapes, potentially compromising their ability to model native cis-regulatory element (CRE) activity. Hepatocyte-like cell lines (HLCs), particularly HepG2, are frequently used to study liver CREs, but whether non-cancer-derived HLCs, such as AML12, better recapitulate native regulatory landscapes remains unclear. Furthermore, cell culture conditions have been shown to influence gene expression and CRE activity, raising the possibility that optimizing culture conditions could improve HLCs’ gene regulatory similarities to primary liver tissue. Together, these questions highlight the importance of characterizing the transcriptional regulatory landscapes of HLCs and evaluating the effects of culture conditions when using these cell lines to study native liver CRE activity.

**Results:** We systematically characterized the molecular features and transcriptional regulatory landscapes of four HLCs and compared them with those of liver tissue. We examined AML12 cells using RT-qPCR and flow cytometry and found that they express some hepatocyte markers but lack key characteristics of mature hepatocytes. We then used ATAC-seq to characterize the chromatin accessibility landscape of AML12. Through transcription factor motif enrichment, functional enrichment, and machine learning analyses, we found substantial differences in the cis-regulatory landscapes of AML12 and primary mouse liver. Modifying AML12 culture conditions altered some regulatory features but did not substantially improve their similarity to primary liver tissue. We then extended these analyses to three additional HLCs and found similar differences between their regulatory landscapes and that of primary liver tissue. Notably, HepG2 recapitulated native liver regulatory landscapes better than the tested non-cancer-derived cell lines.

**Conclusion:** Our findings reveal substantial differences in transcriptional regulatory landscapes between immortalized HLCs and native liver tissue that persist despite modifications to culture conditions. These results highlight the limitations of HLCs in modeling native liver transcriptional regulation and emphasize the importance of evaluating their regulatory landscapes before using them to investigate CRE function.

## Background

Immortalized cell lines are widely used in biological research. They can serve as an inexpensive and experimentally tractable proxy for tissues and cell types that are difficult to obtain or maintain, thereby eliminating the need for animal sacrifice, post-mortem human tissue, or long-term primary cell culture. However, immortalization, prolonged culture, and adaptation to cell culture conditions can cause cell lines to diverge from their tissues or cell types of origin. Such differences have been documented in a variety of cell lines, including endothelial cells(1,2), immune cells(3,4), and neurons(5). In addition, previous studies showed that cancer-derived cell lines are often poor models of their tumors of origin(6–8). These findings can make results from cell line experiments difficult to interpret, particularly when results are used to understand mechanisms of native tissues.

A popular use of immortalized cell lines is for high-throughput assays for deciphering transcriptional regulatory mechanisms. These assays include massively parallel reporter assays (MPRAs)(9–13), in which synthesized DNA sequences are tested for their ability to act as *cis*-regulatory elements (CREs) that activate transcription, and CRISPR screens to dissect CRE-gene relationships(14,15). The ability to assay thousands to tens of thousands of hypotheses regarding CREs in parallel makes these approaches powerful tools for studying regulatory sequence function. However, their interpretation depends on the transcriptional regulatory state of the assayed cells, including transcription factor (TF) activity and chromatin accessibility, making discrepancies between cell lines and their tissues or cell types of origin particularly consequential for studies that use cell lines to make inferences about native CRE activity.

Profiling chromatin accessibility therefore provides a way to assess whether cell lines retain tissue-relevant regulatory environments for interpreting CRE activity. Yet, except for blood and skin(16), comparisons of CRE activity between cell lines and their tissue or cell type of origin have been limited.

The hepatoblastoma-derived cell line HepG2 is one of the most widely used cell lines for CRE activity assays(9–12,17,18). Studies using HepG2 have identified TF motifs–sequences bound by TFs that regulate gene transcription–underlying hepatocyte CRE activity(9,12), tested how sequence differences between primate species(10) and between human individuals(11) influence hepatocyte CRE activity, and provided insights into mechanisms underlying hepatitis B(17) and other hepatocyte-influenced disorders (11). Such results have the potential to provide exciting insights into liver biology, as hepatocytes make up the majority of the liver. However, this potential depends on the extent to which the gene regulatory environment of HepG2 reflects that of primary hepatocytes.

RNA sequencing (RNA-seq) data have indicated that hepatocyte-like cells (HLCs), including HepG2, often lack key hepatocyte characteristics including expression of genes in metabolic pathways, drug-metabolizing enzymes, and transporters; sometimes retain immature phenotypes; and sometimes take on non-liver tissue identities(19–24). At the functional level, lipid accumulation was not observed in response to fructose in multiple HLCs including HepG2(25) and hepatotoxin detection was shown to be limited in multiple HLCs including HepG2(26) despite these being important functions of hepatocytes. Yet, despite these concerns, HLCs are frequently used in liver research(24,27), including in the largest MPRA to date for studying liver CRE activity(12), as maintaining primary hepatocytes in culture for an extended period is infeasible. These concerns highlight the need to assess whether HLCs preserve not only hepatocyte-like gene expression or function but also liver-relevant transcriptional regulatory landscapes.

The studies describing the differences between HLCs and primary hepatocytes focused on HLCs derived from cancer, and cancer is known to alter transcriptional regulatory networks(28–30). Non-cancer-derived HLCs may therefore provide useful alternatives to HepG2 for the study of transcriptional regulatory mechanisms. One example is the alpha mouse liver 12 (AML12) cell line, an immortalized hepatocyte line derived from a healthy transgenic adult mouse by overexpressing human TGFα(31). AML12 cells have become an important model system in liver research, notably in lipid metabolism and liver injury studies(32–35). The study describing the creation of this cell line illustrated that AML12 cells expressed multiple proteins found in mature hepatocytes, including connexins Cx26 and Cx32(36,37), transferrin(38), and Ldh5(39), after four months in culture. Although none of these proteins are expressed exclusively in hepatocytes, their combined expression suggests that AML12 cells are similar to hepatocytes.

In addition, AML12 cells secreted albumin, an important function of hepatocytes, albeit at a lower level than that of primary hepatocytes(31). Notably, they did not express Afp, an immature hepatocyte marker(40–42), or Cx43, a protein expressed in multiple tissue types including nervous system(43) and muscle(44) as well as de-differentiated cell lines and liver bile ducts but not hepatocytes(45), suggesting that they have a mature hepatocyte-specific gene expression program. In addition to AML12 cells, Telomerase-immortalized Liver Epithelial-2 (THLE-2) cells, which come from a healthy human liver, have been used to study how the liver responds to lead(46) and nicotine(47), and Applied Biological Materials (abm) sells non-cancerous hepatocyte-derived cell lines obtained from healthy tissue from five mammalian species. THLE-2 cells have been shown to express ALB and KRT18, which play important roles in hepatocytes, but not AFP(48). Unfortunately, there is little characterization of the HLCs from abm. However, CRE activity in HLCs has not been compared to CRE activity in the liver, limiting our ability to determine the suitability of the cell lines for deciphering transcriptional regulatory mechanisms.

Cell culture conditions have also been shown to have major effects on cell line gene expression, CRE activity, and behavior. For example, recent studies have shown that growing cell lines, including HepG2, in antibiotics can alter gene expression and CRE activity(49,50). Another study found that growing AML12 and THLE-2 cells in media without insulin increases their insulin responsiveness, which was limited under standard cell culture conditions that include supplemental insulin(51). Thus, the utility of MPRAs and other cell line-based assays for studying transcriptional regulatory mechanisms may be dependent not only on the cell line used but also on cell culture conditions.

To investigate the utility of HLCs for studying mechanisms underlying CRE activity, we characterized the transcriptional regulatory regime of four hepatocyte-derived cell lines. First, we analyzed AML12 in multiple culture conditions using RT-qPCR, flow cytometry, and the Assay for Transposase-Accessible Chromatin with sequencing (ATAC-seq), a genome-wide assay of chromatin accessibility that identifies candidate CREs. We compared the results to those from primary liver, pancreas, and cortex tissue from healthy adult mice. Upon finding that AML12 cells grown in all culture conditions have major transcriptional regulatory differences from primary mouse liver tissue, we extended our comparison by performing ATAC-seq on rat (T0078) and human (T0063) hepatocytes from abm as well as HepG2. Although all cell lines had some transcriptional regulatory network similarity to primary liver tissue, they all had major differences, demonstrating the value of profiling CRE activity in cell lines when selecting a system to use to test specific hypotheses regarding transcriptional regulation.

## Methods

### Cell culture

AML12 cells: A frozen vial of 1 million AML12 cells was obtained from ATCC (Cat. No. CRL-2254, Batch No. 70061353) and thawed into Dulbecco’s Modified Eagle Medium/Nutrient Mixture F-12 (DMEM-F12) media (Gibco, Cat. No. 11-320-033) supplemented with 1% 100x Insulin/transferrin/selenium (ITS) (Gibco, Cat. No. 41-400-045), 10% fetal bovine serum (Gibco A5670701), 39 ng/mL dexamethasone (Sigma-Aldrich, Cat. No. D2915) and 100 U/mL penicillin/streptomycin (P/S) (Gibco, Cat. No. 15-140-122). During cell culture, the cells reached approximately 70-90% confluency before being dissociated with 0.25% trypsin (Gibco, 25-200-114) and passaged. For some experiments, AML12s were cultured without insulin for 36-48 hours, substituting the ITS solution for 5.5 μg/mL transferrin (MilliporeSigma, 61624) and 6.7 ng/mL sodium selenite (MP Biomedicals, 0219474110). The transferrin and sodium selenite concentrations were the same as in the final dilution of the Gibco ITS solution. For some experiments, cells were cultured for three weeks without P/S. Cells were passaged 2-3 times for all experiments, except for the samples that were cultured for 3 weeks without P/S, which were passaged 6 times.

abm-human cells: A frozen vial of 1 million immortalized human hepatocytes (Cat. No. T0063) were obtained by Applied Biological Materials (abm) and thawed into PriGrow IX medium (abm, Cat. No. TM019) supplemented with 10% FBS. P/S was not included. The cells were cultured on PriCoat flasks (abm, Cat. No. G299). The cells reached approximately 70-90% confluency before being dissociated with 0.25% trypsin (Gibco, Cat. No. 25-200-114) and passaged. Before ATAC-seq experiments, the cells were transferred to 6-well plates pre-coated with extracellular matrix (ECM) (abm, Cat. No. G422) according to the manufacturer’s protocol. Cells were passaged 2-3 times for all experiments.

abm-rat cells: A vial of 1 million immortalized rat hepatocytes (Cat. No. T0078) were obtained from Applied Biological Materials (abm). The cell culture conditions were kept the same as those for the human hepatocytes from abm. Likewise, P/S was not included. Cells were passaged 2-3 times for all experiments.

HepG2 cells: A frozen vial of 1 million HepG2 hepatocytes was obtained from ATCC (Cat. No. HB8065) and thawed into DMEM (Gibco, Cat. No. 11-995-073). Note that, unlike in previous genomics assays for HepG2(52–54), cells were cultured without P/S. The cells reached approximately 70-90% confluency before being dissociated with 0.25% trypsin (Gibco, Cat. No. 25-200-114) and passaged. Cells were passaged 2-3 times for all experiments.

### Mouse Euthanasia

Mice were sacrificed via isoflurane overdose until deep gasping breaths (agonal breathing) and lack of the pedal withdrawal reflex were observed. At this stage, the mice were immediately decapitated and dissected. The procedure was carried out by A.R. Brown under IACUC procedure #PROTO201600003.

### RNA Extraction and RT-qPCR

RNA extraction from AML12 cells: Prior to RNA extractions, AML12 cells were thawed and cultured in complete media for 2 days before being passaged. 1.1 million cells were seeded in 2 different T75 flasks and incubated overnight before insulin-media was added to two of the flasks so that RNA could be extracted from cells grown in either complete media or cells grown in insulin-media. The cells were incubated for 36 hours, reaching 80-90% confluency before they were dissociated with 0.25% trypsin (Gibco, Cat. No. 25-200-114), transferred to 15 mL conical tubes, counted, washed with ice cold PBS (Gibco, Cat. No. 14-190-250), and pelleted via centrifugation at 4 °C at 500 x g for 5 minutes. All work was completed in an RNAse-free Grant-bio UVT-S-AR hood, and the samples were kept on ice for each step unless otherwise specified. RNA was extracted using the RNeasy Mini Kit (Qiagen, Cat. No. 74104) with a slightly modified version of the manufacturer’s protocol: 600 μL of buffer RLT supplemented with 1% Beta-Mercaptoethanol (Sigma-Aldrich, Cat. No. M3148) was used to resuspend each cell pellet and lyse the cells. The lysate was added to Qiagen QIAshredder tubes (Cat No. 79654) and centrifuged at 13,000 RPM. 600 μL of 70% molecular biology grade ethanol (Decon Labs, Cat. No. 07-678-003) was then mixed into the eluent. The eluent mixture was then passed through a Qiagen RNeasy spin column by centrifuging at 10,000 RPM for 1 minute, and the flowthrough was discarded. The RNeasy column was washed with 350 μL RW1 buffer and centrifuged again at 10,000 RPM for 1 minute. A mixture of 10 μL DNAse I (Qiagen RNase-Free DNase Set, Cat No. 79254) and 70 μL buffer RDD was then added to each sample and incubated for 15 minutes at room temperature. 350 μL RW1 buffer was added to the column, before centrifuging again at 10,000 RPM for 1 minute. The column was then washed with 500 μL of buffer RPE and centrifuged at 10,000 RPM for 1 minute. This step was repeated, and then the column was transferred to a new collection tube and centrifuged again at 13,000 RPM for 1 minute to remove any residual buffer. The column was then transferred to a clean RNAse-free 1.5 mL Eppendorf tube, and the RNA was eluted with 40 μL of RNase free water by centrifuging at 10,000 RPM for 1 minute.

RNA extraction from flash frozen mouse liver: Fresh liver tissue was obtained from A. Brown and placed in a snap-cap 1.5 mL tube. The tube was immediately immersed in liquid nitrogen for 20 minutes, transferred to dry ice, and kept there for 15 minutes to allow residual liquid nitrogen to evaporate. The sample was then transferred to −80°C for 2 months. A pea-sized section of the tissue was immersed in 400 μL ice cold buffer RLT supplemented with 1% Beta-Mercaptoethanol in a 1.5 mL Eppendorf tube for less than 1 minute to thaw and then placed on a plastic petri dish on ice. The sample was minced with a razor blade and returned to the Eppendorf tube with the ice cold RLT buffer. The tissue was further homogenized by passing it through a 20 gauge needle attached to a 1 mL RNase-free syringe 10 times. The inside of the syringe was washed with an additional 300 μL ice cold buffer RLT before the homogenate was transferred to a QiaShredder column. At this stage, the RNA extraction protocol described above for the AML12 cells was used.

RNA quantification and quality control: RNA concentration was quantified with a Qubit™ RNA broad range assay kit (Invitrogen, Cat. No. Q10211) on an Invitrogen Qubit™ 3.0 Fluorometer following the manufacturer’s protocol. The quality of the extracted RNA was assessed using the RNA TapeStation kit (Agilent Cat. No. 5067-5576 and Cat. No. 5067-5577) on the Agilent 4150 TapeStation system following the manufacturer’s protocol. The quality of the RNA was assessed through the RNA Integrity Number (RIN) calculated by the TapeStation analysis software. The RINs of all samples were at least 8.0 **(Table S1)**, indicating that high-quality RNA was obtained.

RNA to cDNA conversion: The RNA was converted to cDNA using the Applied Biosystems High-Capacity RNA-to-cDNA™ Kit (Cat No. 4387406) following the manufacturer’s protocol. The concentration of the resulting cDNA was then quantified with an Invitrogen™ Qubit™ 1X dsDNA High Sensitivity Kit ( Cat No. Q33266).

RT-qPCR: RT-qPCR experiments were performed using TaqMan gene expression assays with probes testing the relative expression levels of the following genes of interest: Ctcf (Mm00484027_m1), Gapdh (Mm99999915_g1), Dbx1 (Mm02344179_m1), Hnf4A (Mm00433959_m1), Onecut1 (Mm00839394_m1), Alb (Mm00802090_m1), Cd68 (Mm03047343_m1), and B2m (Mm00437762_m1). First, the concentration of the cDNA was adjusted to 10 ng/μL through dilution with nuclease-free water. Then, 10 μL Taqman advanced master mix (Applied Biosystems, Cat. No. 4444963), 1 μL Taqman probe, 5 μL PCR grade water, and 2 μL cDNA were combined in each well of a 96-well optical grade qPCR plate (Applied Biosystems, Cat. No. N8010560). The plate was sealed with MicroAmp™ Optical Adhesive Film (Applied Biosystems, Cat. No. 4311971) and loaded onto a QuantStudio3 qPCR system using the following program: UNG incubation: 50 °C, 2 minutes 1x; polymerase activation: 95 °C, 20 seconds 1x, (denature, 95 °C, 1 sec, Anneal/extend 60 °C, 20 seconds) x 40. The results were then analyzed using QuantStudio Design and Analysis Software v1.6.1. The Ct value from each amplified gene was then normalized by subtracting the Ct value of the housekeeping gene B2m.

### Flow cytometry

AML12 cells were thawed and cultured in complete media for 8 days prior to conducting flow cytometry experiments. The cells were dissociated from the cell culture flasks with 0.25% trypsin before centrifuging at 1500 RPM for 5 minutes and resuspending them in PBS supplemented with 2% fetal bovine serum. Cells count and viability were assessed using a Countess II FL automated cell counter (Invitrogen). Both cell lines showed viability >95%. Vials of 10 million cells each were resuspended to 500 μL and stained with 1:1000 LIVE/DEAD™ Fixable Violet, (Thermo, Cat. No. L34963) for 10 minutes. They were then washed with PBS supplemented with 2% FBS before being stained with 250 μL of 1:100 APC anti-mouse CD24 Recombinant Antibody (Biolegend Cat. No. 114303), 1:100 PE anti-mouse CD68 (Biolegend Cat. No. 137013), and 1:80 ASGR1 Polyclonal Antibody, CoraLite® Plus 488 (Thermo, Cat. No. CL488-11739). Staining proceeded for 20 minutes in the dark at room temperature. Each sample was washed and then resuspended in 3 mL PBS supplemented with 2% FBS. Large clumps of cells were removed by passing the cells through a 35 μm Nylon Mesh Screen into 5 mL FACS tubes (Falcon, Cat No. 352235). Flow cytometry was carried out on a Becton Dickinson FACSVantage SE with FACSDiva option.

### ATAC-seq

ATAC-seq was executed on AML12, abm-rat, abm-human, and HepG2 cells as described in previous work(55–58) with some modifications.

Sample preparation and tagmentation: All cell lines were cultured as described above. 250,000 cells were plated in 6-well tissue culture treated plates (Starstedt, Cat. No. 83.3920). Each cell culture condition tested was carried out in duplicate for all cell lines. The cells were grown for 36-48 hours before being washed with ice cold PBS. Nuclei were extracted by adding 2 mL ice cold lysis buffer (15 mM Tris-HCl, pH 7.5; 15 mM NaCl; 4.5 mM MgCl2; and 10% IGEPAL in MilliQ water). The cells were incubated in the lysis buffer for 5-10 minutes or until the cell membranes became visibly porous under a phase contrast microscope. The lysed cells were then dissociated from the surface of the cell culture plate through gentle pipetting, and the suspension was collected in 15 mL conical tubes. The tubes were centrifuged at 500 x g for 5 minutes at 4 °C, causing the intact nuclei to form a pellet. The nuclei were resuspended in 50-200 μL nuclease-free water. An aliquot of the nuclei was stained with DAPI (Invitrogen, Cat. No. 62248) and counted using a Countess II FL automated cell counter. The gating was adjusted so that the countess correctly identified nuclei and an accurate nuclei count was obtained.

The tagmentation reaction was carried out using an Illumina Tagment DNA Enzyme and Buffer Kit (Cat. No. 20034197). 25 μL of buffer and 2.5 μL Tn5 transposase were combined in a 1.5 mL Eppendorf tube. 50,000 nuclei were then added to the mixture (variable volume), and the total reaction volume was brought to 50 μL with nuclease-free water. The transposition reaction was then incubated on an Eppendorf ThermoMixer C heat block at 37 °C while shaking at 300 RPM for 30 minutes. The reaction was immediately cooled on ice before the transposed DNA was purified using a Qiagen MiniElute PCR Purification Kit (Cat. No. 28004) following the manufacturer’s protocol.

Amplification and size selection of transposed DNA: Each sample was amplified with PCR in a 50μL reaction volume containing 25 μL NEBNext High-Fidelity 2x PCR Master Mix (NEB, Cat. No. M0541),10 μL transposed DNA, 12.5 μL of 10 μM forward and reverse primers that contained a unique combination of Nextera index adaptor sequences, and 2.5 μL nuclease free water. The transposed DNA was amplified using the following PCR program: (5 minutes, 72 °C

→ 30 seconds, 98 °C)x1 → (10 seconds, 98 °C → 30 seconds 63 °C → 1 minute 72 °C)x5.

The PCR amplification was then paused to perform a qPCR side reaction so that the optimal number of additional amplification cycles could be calculated for each sample. The qPCR reactions were set up as follows: 5 μL of previously amplified DNA, 1.25 μL primer mix, 5 μL NEBNext High-Fidelity 2x PCR Master Mix, 0.95 μL 10x SYBR Green, and 2.80 μL nuclease-free water. The following qPCR amplification program was run on a QuantStudio3 qPCR system: (30 seconds, 98 °C) x1 → (10 seconds, 98 °C → 30 seconds 63 °C → 1 minute 72 °C

→ capture fluorescence) x20. The cycle number where the fluorescence attains ⅓ of its maximum intensity corresponds to the optimal number of additional PCR amplification cycles that a given sample needs. The amplification of the transposed DNA is completed with the following PCR program: (30 seconds, 98 °C) x1 → (10 seconds, 98 °C → 30 seconds 63 °C → 1 minute 72 °C) x N, where N is the computed optimal number of cycles.

AMPure XP Beads for DNA Cleanup (Beckman-Coulter, Cat. No. A63880) were used to select for the desired fragment length of approximately 100-1000 base pairs. 40 μL of the PCR-amplified transposed DNA was gently mixed with 22 μL of beads in a PCR tube, incubated at room temperature for 5 minutes, and then separated on a magnetic stand to remove the beads and large DNA fragments bound to them. Leaving the tubes on the stand, 62 μL of supernatant was collected, and combined with 50 μL of fresh beads. They were incubated at room temperature for 5 minutes before being returned to the magnetic stand to remove the beads from the suspension. This time, the supernatant was discarded, as the desired DNA fragments are bound to the beads. The beads were left on the magnetic stand and washed twice with 200 μL of 70% molecular biology grade ethanol before drying fully for 15 minutes. The beads were then resuspended in 12 μL EB buffer (Qiagen, Cat. No. 19086). The beads were separated on the magnetic stand, and 10 μL supernatant was collected for QC and deep sequencing.

DNA quantification and QC: DNA concentration was quantified with a Qubit™ dsDNA high sensitivity assay kit (Invitrogen, Cat. No. Q32854) on an Invitrogen Qubit™ 3.0 Fluorometer following the manufacturer’s protocol. The quality of the extracted DNA was assessed using the D1000 DNA TapeStation kit (Agilent Cat. No. 5067-5582 and Cat. No. 5067-5602) on an Agilent 4150 TapeStation system according to the manufacturer’s protocol. The quality of the transposed DNA library was assessed qualitatively by analyzing the periodicity of the TapeStation plot (**Figure S1**) on the TapeStation analysis software; all samples with the expected periodicity(55) were pooled and sent for sequencing.

Sequencing: Samples were sequenced by Azenta (GeneWiz) through their next generation sequencing (NGS) service. Samples were run on the NovaSeq X Plus sequencing system using the 2x 150 bp sequencing mode. Replicates were not split across runs.

### ATAC-seq data processing

ATAC-seq data generated in this study were processed using the ENCODE ATAC-seq pipeline (v2.2.3)(59), starting from raw FASTQ files. Default pipeline parameters were used except that automatic adapter detection was enabled (“atac.auto_detect_adapter”: true) and only uniquely mapped reads were retained (“atac.multimapping”: 0). Reads from *Homo sapiens* (human) were mapped to hg38, reads from *Mus musculus* (mouse) were mapped to mm10, and reads from *Rattus norvegicus* (rat) were mapped to rn7. Data quality reports can be found at https://github.com/KaplowLab/Comparison_of_liver_cell_line_and_native_liver and 10.5281/zenodo.20544805.

Human liver tissue ATAC-seq data were obtained from ENCODE (ENCSR952SPO, ENCSR685ZMP) and GEO (GSM5021293, GSM5021292)(60). The data were selected based on either the reported health status of the sample donors or the TSS enrichment scores available before data reprocessing. Specifically, ENCSR952SPO and ENCSR685ZMP were included because their donors were not recorded as having any disease diagnoses, unlike other human adult liver samples available from ENCODE. GSM5021293 and GSM5021292 were selected because they had relatively high TSS enrichment scores compared with other samples in the same dataset (60), and their donors were also not recorded as having liver-related disease diagnoses. The two PHH ATAC-seq datasets were obtained from GSE185360 and GSE207590 (GSM6300669, GSM6300670)(61,62). Human pancreas ATAC-seq data were obtained from ENCODE (ENCSR705KEB, ENCSR918TVE, ENCSR808ZMK, ENCSR530XBF, ENCSR251POP)(52). Human cortex ATAC-seq data were obtained from primary motor cortex neurons from (63). Adult mouse liver tissue ATAC-seq data were obtained from (57,64).

Newborn mouse liver tissue was obtained from ENCODE (ENCSR609OHJ)(52). Mouse pancreas tissue ATAC-seq data were obtained from (65). Mouse cortex ATAC-seq data were obtained from (66). Adult rat liver tissue ATAC-seq data were obtained from (67,68).

All ENCODE datasets were processed using the ENCODE ATAC-seq pipeline (V2.2.3) starting from the filtered BAM files; the two PHH datasets were processed starting with the FASTQ files, and the human non-brain as well as the mouse tissue samples were processed as described in our previous work(57). The parameters and genome assemblies described above were used for all data processing.

### Principal Component Analysis (PCA)

All ATAC-seq peak sets were pooled by species to generate consensus peak sets for PCA analysis. Human-derived samples included bulk liver, PHH, abm-human, HepG2, pancreas, and cortex. Mouse-derived samples included adult bulk liver, newborn bulk liver, AML12 under all media conditions, pancreas, and cortex (7 weeks and 12 weeks). Chromatin accessibility was quantified from filtered BAM files over the corresponding consensus peak sets using featureCounts(69), generating count matrices containing all sample replicates. These count matrices were subsequently used for PCA.

### Motif enrichment analysis

Motif enrichment analysis was performed using MEME-CHIP(70). IDR (Irreproducible Discovery Rate) optimal peaks were used when more than two replicates were available; otherwise, IDR conservative peaks were used. Enhancers were defined as regions that: (1) were located more than 20 kb away from the nearest transcription start site (TSS), (2) were no longer than 1 kb, and (3) did not overlap Coding Sequence (CDS). Human TSS annotations were derived from FANTOM5 hg38 CAGE peaks(71), and CDS annotations were defined using GENCODE v27(72). Mouse TSS annotations were derived from FANTOM5 mm10 CAGE peaks(71), and CDS annotations were defined using GENCODE vM15(72). Rat TSS annotations in rn7 were generated by running liftOver(73) with FANTOM5 rat (rn6) CAGE peaks, and CDS annotations were defined using the rn7 GFF file obtained from https://ftp.ncbi.nlm.nih.gov/genomes/all/GCF/015/227/675/GCF_015227675.2_mRatBN7.2/GCF_015227675.2_mRatBN7.2_genomic.gff.gz(74). The candidate enhancer peaks were centered on the peak summits and resized to a fixed window of 500 bp for motif enrichment analysis.

For motif enrichment analysis of general peak sets, “Classic mode” was used. For motif analysis of DESeq2 peak sets, “Discriminative mode” was used. Discriminative mode was run in both directions, with regions more open in one group used as the primary sequence set and regions more open in the other group used as the control sequence set, and vice versa. Default MEME-CHIP parameters were used except that the number of motifs to be discovered by MEME was set to 10. CIS-BP 2.0 single-species DNA motif databases(75) corresponding to the species being analyzed were used. In human samples, ONECUT1 results were obtained from STREME(76), whereas all remaining motif enrichment results were derived from CENTRIMO(77). In mouse samples, ONECUT1 results for mouse liver from Liu *et al*.(65) and FOXA1 results for AML12 grown in complete media were obtained from STREME, whereas all remaining results were derived from CENTRIMO. In rat samples, all motif enrichment results were derived from CENTRIMO.

### Gene Ontology enrichment analysis

rGREAT(78) was used to identify biological processes enriched among genes associated with peaks and differentially open peaks. Enrichment analysis was performed using the ‘great’ analysis function across three Gene Ontology (GO) categories: Biological Process (BP), Cellular Component (CC), and Molecular Function (MF). Results from all ontologies were combined and filtered to retain terms with fold enrichment ≥ 2 and binomial adjusted p-value < 0.05.

### Machine learning model evaluation

The multi-species liver enhancer classification model trained on primary liver data from *Mus musculus*, *Rattus norvegicus*, *Macaca mulatta*, *Bos taurus*, and *Sus scrofa* presented in our previous work(79) was used for all machine learning model evaluations. Although we did not evaluate this model using human data in our previous work, in this study, we found high sensitivity in multiple human liver datasets (**Figure 4C**). For each evaluation, only enhancers were used because only enhancers were used to train the model. Models were evaluated on only enhancers on mouse chromosomes 1 and 2, which were the test set chromosomes in our previous work(57,79), to prevent evaluation on enhancers that might have been used in training or hyper-parameter tuning. When evaluating the model on the intersection of AML12 enhancers, AML12 enhancers from complete media that overlapped AML12 enhancers from every other condition were used; such enhancers were found using bedtools version 2.29.9 intersect with options -wa and -u(80). When evaluating the model on the intersection of mouse liver enhancers, the mouse test set from our previous work(57,79) was used.

For enhancers from rat and human datasets, test set enhancers, which were those that mapped to mouse test chromosomes, were used, as described in our previous work(57,79). Specifically, rat enhancers were mapped from the rn7 assembly to the rn6 assembly with liftOver(73), and then HALPER(81) was run with options -max_frac 1.5, -min_len 50, -protect_dist 5, and - narrowPeak to remove enhancers with very different lengths between assemblies. Rat enhancers were next mapped to the house mouse mm10 assembly using halLiftover version 2.1(82) with the Zoonomia Cactus alignment(83,84), and then HALPER was run with the same settings as before to create contiguous orthologs and remove those with very different lengths in different species. Human enhancers were mapped from the hg38 assembly to the mouse mm10 assembly in the same way that rat enhancers were mapped to the mm10 assembly. After mapping to the mm10 assembly, enhancers on mouse chromosomes 1 and 2 were selected, and their peak names were used to filter the original rat and human enhancer files for those on the test set chromosomes. The intersection of rat enhancers across datasets from fresh(67) and flash-frozen(68) tissue was found using bedtools version 2.29.9 intersect with options -wa and - u(80), where -a was the enhancers from fresh tissue and -b was the enhancers from flash-frozen tissue. The intersection of human enhancers across data from cryo-preserved(52) and flash-frozen(60) tissue was found using bedtools version 2.29.9 intersect with the same options(80), where -a was the enhancers from cryo-preserved tissue and -b was the enhancers from flash-frozen tissue.

### Differential chromatin accessibility analysis

Differential chromatin accessibility analysis was performed using DESeq2(85) on count matrices generated from the consensus peak sets described in the above “Principal Component Analysis (PCA)” section. DESeq2 accounts for differences in read depth between samples(85); thus, it should not identify peaks solely explained by differences in sequencing depths between the cell line and primary liver tissue datasets. Human comparisons included abm-human, HepG2, human bulk liver, and PHH samples. Mouse comparisons included AML12 cells grown under all media conditions and both adult mouse bulk liver datasets. Rat comparisons included abm-rat and both adult rat bulk liver datasets. For each pairwise comparison, raw read counts within peak regions were analyzed using DESeq2 with default settings for size-factor normalization and dispersion estimation. Peaks were classified as more open or less open based on the sign of the log2 fold change. Statistically significant was defined as an adjusted p-value (Benjamini-Hochberg FDR) < 0.05, and differential accessibility was further required to have an absolute log2 fold change ≥ 1.

## Results

### Only a strict subset of mature hepatocyte marker genes are expressed in AML12 cells

To assess the extent to which AML12 cells are similar to primary liver tissue, we evaluated the expression of genes with important roles in the liver. We extracted RNA from AML12 cells cultured in complete and insulin-deficient media as well as from primary mouse liver tissue. We then used reverse transcription quantitative polymerase chain reaction (RT-qPCR) to quantify gene expression (**Figure 1A**). We assayed the expression of Ctcf and Gapdh, which are expressed across all adult mouse tissues in which they have been assayed(86) and thus served as positive controls; Dbx1, which is expressed in the developing brain and not the liver and thus served as a negative control; Hnf4a(87), Onecut1(88), and Alb, which are expressed in the liver but not many other tissues; and Cd68(89), which is expressed in Kupffer cells, a minority liver cell type, but not hepatocytes. We normalized the expression levels of these genes to the housekeeping gene B2m(90) and computed the ΔCt value. Relative to B2m, AML12 cells expressed the known hepatocyte TFs Hnf4a and Onecut1 as well as Ctcf and Gapdh at similar levels to bulk mouse liver. Alb expression in AML12 cells was lower than in mouse liver, as was previously reported(31). We detected trace levels of Dbx1 in AML12 cells but no Dbx1 expression in bulk mouse liver. Unexpectedly, the AML12 cells and mouse liver expressed similar levels of Cd68 even though AML12 cells should not contain any non-hepatocyte liver cells(31). These results were consistent for AML12 cells grown in complete media and insulin-deficient (-insulin) media. While the detection of Onecut1, Hnf4a, and Alb illustrates that AML12 cells express some key hepatocyte markers, the lower expression of Alb suggests that AML12 cells might not be fully mature, and the detection of high levels of Cd68 suggests that they may harbor a previously undetected population of Kupffer cells or express genes not typically associated with hepatocytes.

**Figure 1.**
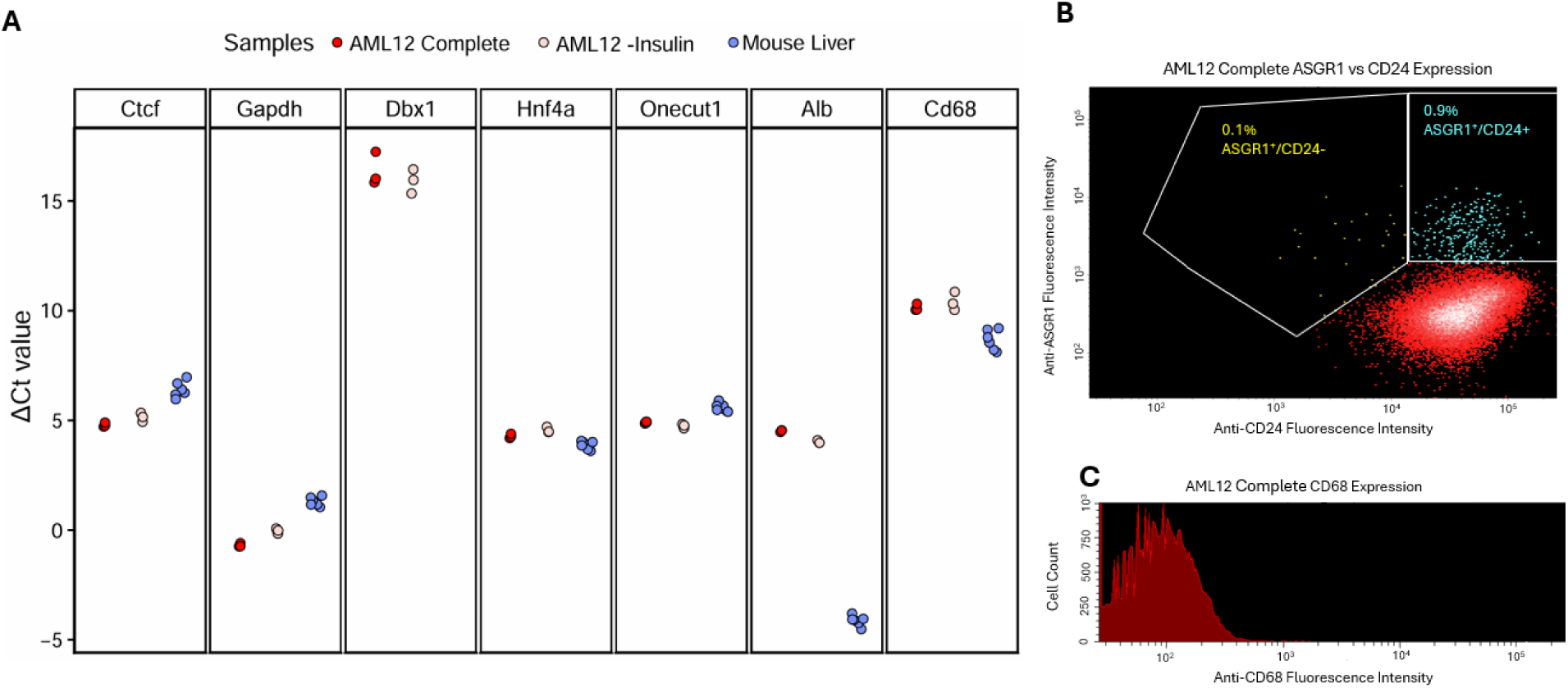
AML12 cell RT-qPCR and flow cytometry provide conflicting information on the extent of similarity between AML12 cells and mature hepatocytes. A) RT-qPCR of AML12 cells cultured with or without insulin and compared to bulk mouse liver. Lower ΔCt values indicate higher gene expression. B) Flow cytometry analysis of AML12 cells grown in complete media. The fluorescence intensity of CD24 (x-axis) vs ASGR1 (y-axis) is used to identify the proportion of cells that express one or both of these surface markers. C) The fluorescence intensity of CD68 on the x-axis plotted against cell count on the y-axis. The cells in both panels were triple-stained with anti-CD24-APC, anti-ASGR1-CorralitePlus488, and anti-CD68-PE and have been gated through forward/side scatter to eliminate auto-fluorescence of dead cells and debris.

We then used flow cytometry to determine if AML12 cells constitute a population of pure HLCs or if there might be a mixture of hepatocytes and other cell types such as Kupffer cells. We sorted AML12 cells based on surface protein levels of ASGR1, a mature hepatocyte marker(91,92); CD24, which is on the surfaces of immature hepatocyte precursor cells(93) as well as some other cell types in the hematopoietic lineage; and CD68, which is on the surface of activated Kupffer cells(94). Flow cytometry analysis indicated that over 99% of the population is positive for CD24. In addition, 0.9% of the population expresses both ASGR1 and CD24, and only 0.1% express ASGR1 alone (**Figure 1B**). Furthermore, flow cytometry analysis indicated that AML12 cells are uniformly negative for surface expression of CD68 (**Figure 1C**). Taken together, this indicates that AML12 cells probably do not harbor a distinct population of previously undetected Kupffer cells but lack some key characteristics of mature hepatocytes while expressing CD24 instead.

### AML12 cells have a different transcriptional regulatory landscape from primary liver tissue

This discrepancy between the high expression of TFs that control hepatocyte transcription in AML12 cells indicated by the RT-qPCR results and the lack of AML12 cell hepatocyte identity indicated by the flow cytometry results raised the question of whether AML12 cells retain a liver-like transcriptional regulatory landscape. To address this question, we performed ATAC-seq(55) on AML12 cells and compared the results to published mouse liver datasets(57,65). ATAC-seq identifies accessible chromatin regions, which can be considered candidate CREs because they are available for TF binding, allowing us to directly assess the extent to which AML12 cells recapitulate the transcriptional regulatory landscape of primary liver tissue.

TF motif(70,75) and Gene Ontology (GO)(78,95) enrichment analyses showed that AML12 cells retain limited hepatocyte-associated regulatory features. For motif enrichment analysis, we focused on distal candidate CREs, which we refer to as enhancers, because enhancer activity is often highly tissue- and cell-type-specific relative to promoter activity(96). This focus allowed us to better capture TF motifs associated with tissue-specific transcriptional regulation. AML12 enhancers showed enrichment for motifs of the canonical hepatocyte-associated TFs HNF4A and FOXA1(87,97,98), consistent with the RT-qPCR detection of Hnf4a. However, HNF4A enrichment was much weaker than in mouse liver enhancers (E-value = 5.70E-17 in AML12, E-value = 9.90E-188 and 4.70E-249 in the two mouse liver datasets), and CEBPA enrichment was absent in AML12 enhancers despite its known role in the liver(97,99) and its strong enrichment in enhancers from both mouse liver datasets (E-value = 2.50E-128 and 1.40E-172; **Figure 2A**, **Tables S2-S4**). Likewise, GO analysis of genes near AML12 CREs showed some mitochondria-related functions potentially linked to the liver, but most of the enriched processes were not specific to hepatocytes. These processes included podosome organization, rRNA binding, and unfolded protein binding. In contrast, mouse liver CREs were near genes that tended to be involved in liver metabolic functions including gluconeogenesis, detoxification, lipid homeostasis, cholesterol efflux, and plasma lipoprotein particle organization (**Table S5**).

**Figure 2.**
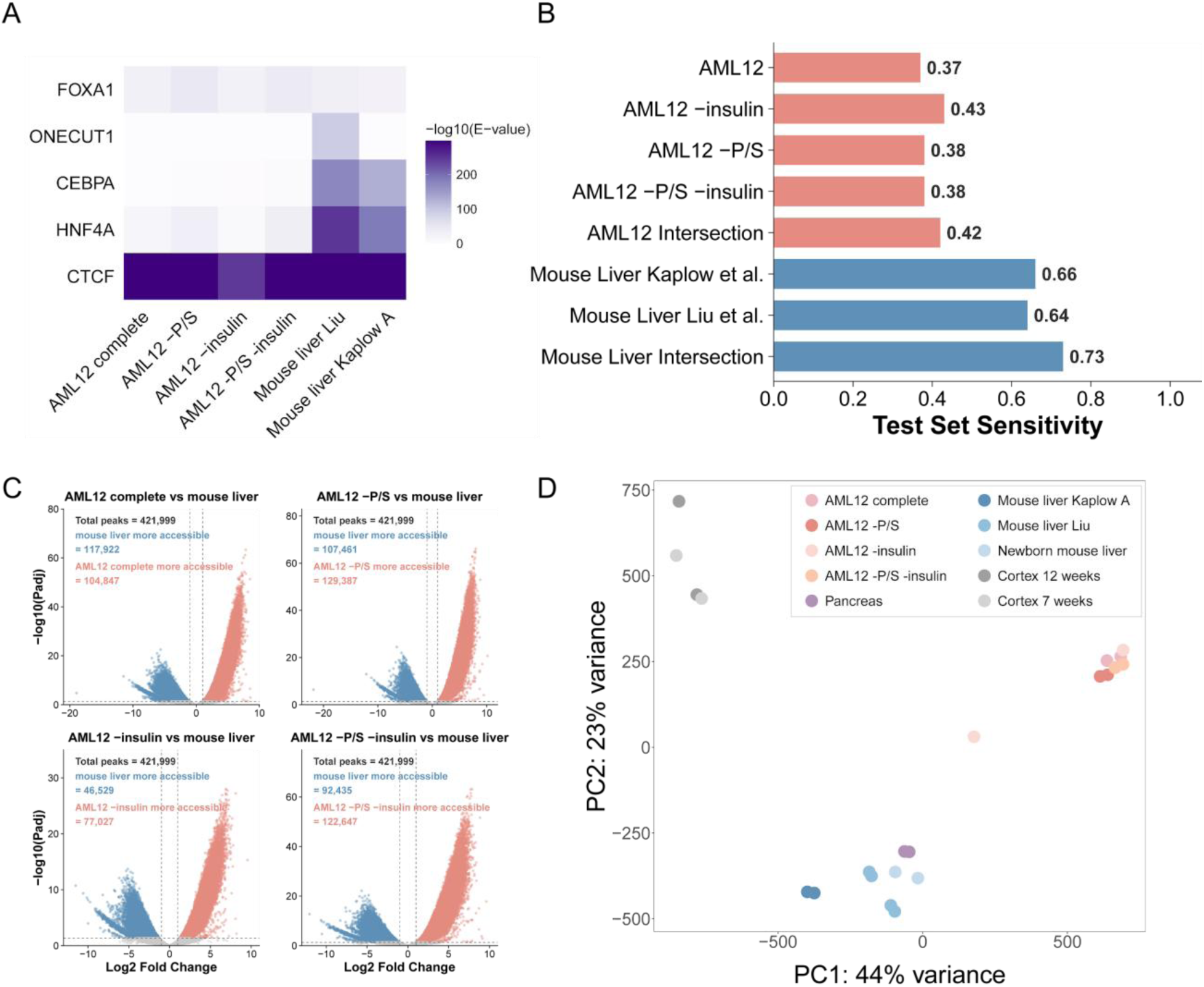
Media modification does not restore a mouse liver-like regulatory landscape in AML12 cells. **A)** MEME-ChIP motif enrichment heatmap of enhancers from AML12 culture conditions and mouse liver datasets. Color indicates motif enrichment significance as -log10(E-value), where motifs that were not detected were assigned an E-value of 1. **B)** Sensitivity of liver machine learning model on mouse enhancers. **C)** DESeq2 differential accessibility volcano plots comparing ATAC-seq peaks between mouse AML12 cell line and reference liver samples. Each point represents an accessible chromatin peak from comparisons of AML12 against bulk mouse liver. The x-axis shows log2 fold change in accessibility, and the y-axis shows -log10 adjusted p-value. Peaks with adjusted p < 0.05 and |log2 fold change| > 1 are highlighted, with red indicating peaks more accessible in cell lines and blue indicating peaks more accessible in liver. **D)** Principal component analysis (PCA) of ATAC-seq from AML12 cells grown in complete medium, without P/S (-P/S), without insulin (-insulin), or without both P/S and insulin (-P/S - insulin) compared with adult and newborn mouse liver datasets as well as adult cortex and pancreas datasets.

To further investigate the extent to which the TF motifs and other sequence patterns governing AML12 cells’ enhancer activity are similar to those of primary liver tissue, we evaluated the sensitivity of previously published machine learning models trained for predicting liver enhancer activity across species(79) on AML12 cell enhancers. High sensitivity–consistently predicting that AML12 cell enhancers are enhancers–would indicate that enhancers in AML12 cells have sequence patterns that are indicative of liver enhancer activity. Unfortunately, the sensitivity was low, with less than half of AML12 enhancers being predicted as enhancers by the model (**Figure 2B**), demonstrating that, not only do AML12 cells have many differences in enhancer activity from mouse liver, but these differences are associated with differences in sequence patterns found in the enhancers.

Differential accessibility analysis between AML12 complete-media cells and mouse liver further revealed widespread transcriptional regulatory divergence (**Figures 2C-D**). Motif enrichment analysis of differentially accessible enhancers showed that sequences of enhancers more accessible in mouse liver were enriched for HNF4A and CEBPA motifs, whereas sequences of enhancers more accessible in AML12 cells lacked enrichment for these canonical hepatocyte-associated TF motifs (**Tables S6-S7**). Likewise, CREs more accessible in mouse liver were enriched for occurring near genes in biological processes related to metabolic and transport functions, including lipid and cholesterol metabolism, amino acid catabolism, and carbohydrate metabolism. In contrast, genes near CREs more accessible in AML12 cells were enriched for occurring in biological processes less associated with hepatocytes, such as cytoskeletal organization, cell adhesion, apoptosis, and stress-related processes (**Table S8**). Together, these results indicate that AML12 cells’ chromatin accessibility landscape does not recapitulate the motifs or functional programs characteristic of mouse liver.

Given the substantial divergence between AML12 cells grown in complete media and mouse liver tissue, we next evaluated whether altering culture conditions could improve AML12 cells’ hepatocyte-like regulatory features. We focused on penicillin/streptomycin (P/S) and insulin because both components have been reported to affect cell state or gene expression in culture. P/S is an antibiotic mixture commonly used to prevent contamination and was included in our AML12 complete culture medium. However, previous studies have shown that antibiotics such as P/S can influence gene expression profiles in cultured cell lines(49). In addition, AML12 insulin response mechanisms may be altered when cells are grown in insulin-containing media(51). We therefore performed ATAC-seq on AML12 cells grown without P/S (-P/S) for three weeks, without insulin for 36 hours (-insulin), and without both P/S and insulin (-P/S -insulin), and compared these cells with AML12 cells grown in complete media and with mouse liver tissue.

Global chromatin accessibility patterns indicated that media modifications did not make AML12 cells’ transcriptional regulatory landscape more hepatocyte-like. In a principal component analysis (PCA) including AML12 cells grown under different culture conditions, adult and newborn mouse liver, and other mouse tissues, AML12 samples clustered together across conditions rather than with adult mouse liver along the first two principal components (**Figure 2D**). In contrast, newborn mouse liver and mouse pancreas clustered closer to adult mouse liver than any AML12 media condition. Thus, removing insulin, P/S, or both did not substantially shift AML12 cells toward a hepatocyte-like chromatin accessibility landscape. Motif and GO enrichment analyses also indicated that media changes had only limited effects on AML12 transcriptional regulatory signatures. HNF4A motif enrichment was stronger in AML12 -P/S (E-value = 5.40E-35) and -P/S -insulin (E-value = 6.70E-36) cells than in AML12 cells grown in complete media (E-value = 5.70E-17), whereas AML12 -insulin cells showed weaker HNF4A enrichment (E-value = 3.00E-09). Similarly, CEBPA motifs were absent in enhancers from AML12 cells grown in complete media, weakly detectable in AML12 -insulin cells (E-value = 4.30E-02), and more strongly enriched in AML12 -P/S (E-value = 2.90E-03) and -P/S -insulin cells (E-value = 4.30E-07). However, HNF4A and CEBPA motif enrichment across all AML12 cell conditions was substantially weaker than in mouse liver enhancers (**Figure 2A**, **Tables S9-S11**). GO enrichment analysis further showed that genes near CREs in AML12 cells grown in both complete and modified media were consistently associated with biological processes unrelated to hepatocytes (**Table S5**).

Furthermore, the machine learning model did not achieve higher sensitivity (accuracy for enhancers) on enhancers from AML12 cells grown in alternative culture conditions than it did on AML12 cells grown in complete media (**Figure 2B**). Since the model was trained on the intersection of enhancers assayed from fresh tissue(57) and flash-frozen tissue(65,79) and achieved higher sensitivity on enhancers found in both datasets than enhancers found in either dataset (**Figure 2B**), we thought that enhancers found in multiple culture conditions might have a more similar transcriptional regulatory code to liver enhancers than those found in only one condition. We therefore also evaluated the model on the subset of enhancers found in all four conditions, hoping that these shared enhancers would have a similar regulatory code to liver enhancers. Unfortunately, the sensitivity was still poor, with over half of AML12 enhancers predicted as non-liver enhancers (**Figure 2B**).

Comparative analysis of differentially accessible regions further indicated that media changes were insufficient to restore a mature liver transcriptional regulatory landscape in AML12 cells (**Figure 2C**). Across comparisons between mouse liver and AML12 cells grown in alternative media, enhancers more accessible in mouse liver retained strong enrichment for HNF4A and CEBPA motifs. In contrast, enhancers more accessible in AML12 generally lacked enrichment for motifs of key liver TFs including HNF4A(87), CEBPA(99), ONECUT1(88), and FOXA1(98) (**Tables S12-S17**). Genes near CREs more accessible in mouse liver were enriched for involvement in hepatocyte-associated metabolic functions, including cholesterol and sterol metabolism, fatty acid catabolism, reverse cholesterol transport, oleic acid response, and amino acid or pentose phosphate metabolism. In contrast, genes near CREs more accessible in AML12 cells were consistently enriched for GO terms including stress fiber organization, contractile actin filament bundles, podosomes, actomyosin-associated processes, and collagen biosynthetic regulation; these results were consistent across cell culture conditions. CREs more accessible in the AML12 -P/S -insulin condition relative to mouse liver additionally showed enrichment for involvement in neuronal and synaptic signaling (**Table S8**). Taken together, these results indicate that, although removing P/S or insulin altered some components of the transcriptional regulatory landscape, these media modifications were insufficient to resolve most of the transcriptional regulatory divergence between AML12 cells and primary mouse liver tissue.

### Other liver cell lines’ transcriptional regulatory landscapes have major differences from hepatocytes

Because optimizing AML12 culture conditions did not restore a hepatocyte-like regulatory landscape, we next investigated whether other HLCs better preserve hepatocyte regulatory identity. We selected three HLCs with distinct biological origins and immortalization strategies: the Applied Biological Materials T0078 rat hepatocyte cell line (abm-rat), the Applied Biological Materials T0063 human hepatocyte line (abm-human), and HepG2. abm-rat was immortalized through serial passaging and lentiviral transduction with the SV40 large T antigen, whereas abm-human was generated using HPV E6/E7, hTERT, and MycT58A(100). HepG2 was derived from a liver tumor from a 15-year-old that was initially classified as hepatocellular carcinoma(101) but is now considered hepatoblastoma(102). It is one of the most widely used human liver-derived models for studying hepatic gene regulation, although previous work has shown that its chromatin accessibility landscape can differ from that of healthy hepatocytes(30). We therefore used ATAC-seq to evaluate whether any of these cell lines more closely recapitulate the chromatin accessibility landscape of primary hepatocytes or liver tissue.

Comparison between abm-rat and primary rat liver revealed significant loss of hepatocyte-like transcriptional regulatory features in abm-rat. Rat liver enhancers showed strong enrichment for HNF4A (E-value = 8.9e-774 and 6.0e-1611), CEBPA (E-value = 2.00E-129 and 1.2e-418), and FOXA1 (E-value = 4.20E-41 and 7.70E-157) motifs. In contrast, abm-rat enhancers showed only weak HNF4A enrichment (E-value = 4.20E-05) and lacked detectable CEBPA and FOXA1 enrichment (**Figure 3A**, **Tables S18-S20**). Likewise, the sensitivity of the machine learning model for predicting liver enhancer activity(79) was even worse for the abm-rat enhancers than it was for the AML12 enhancers despite strong performance in primary rat liver tissue (**Figure 3B**). This further demonstrates that the abm-rat enhancers’ sequence patterns are often different from those found in primary liver enhancers.

**Figure 3.**
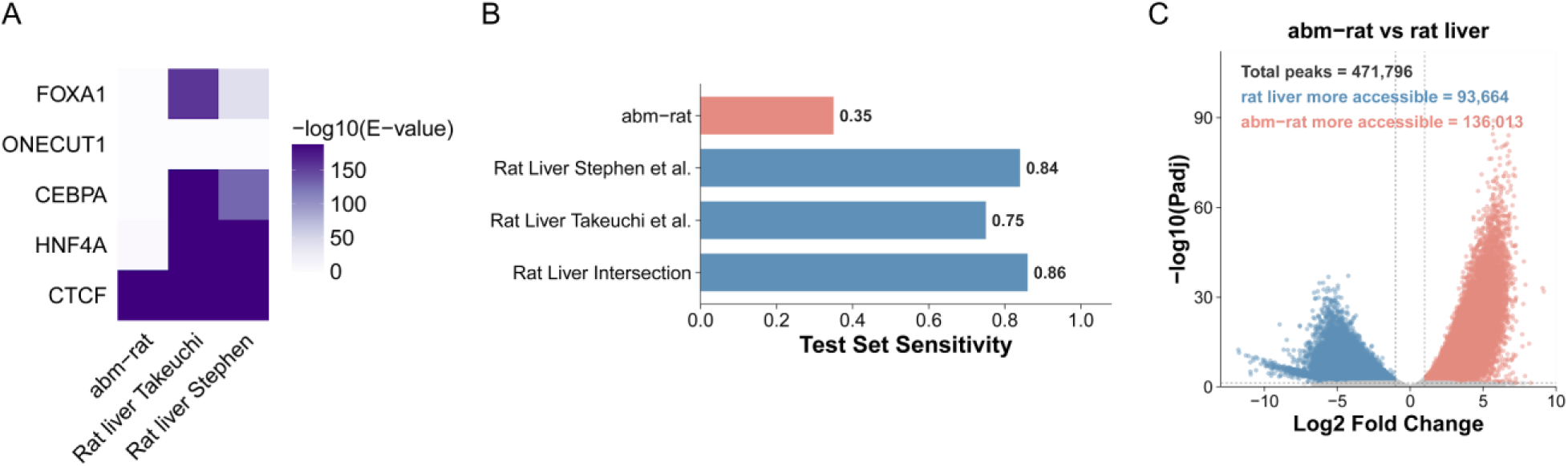
Liver-derived rat cell lines show limited hepatocyte-like regulatory identity. **A)** MEME-ChIP motif enrichment heatmap of enhancers from abm-rat cells and rat liver datasets. Color indicates motif enrichment significance as -log10(E-value), where motifs that were not detected were assigned an E-value of 1. **B)** Sensitivity of liver machine learning model on rat enhancers. **C)** DESeq2 differential accessibility volcano plots comparing ATAC-seq peaks between liver-derived rat cell line and reference liver samples. Each point represents an accessible chromatin peak from comparisons of abm-rat against bulk rat liver. The x-axis shows log2 fold change in accessibility, and the y-axis shows -log10 adjusted p-value. Peaks with adjusted p < 0.05 and |log2 fold change| > 1 are highlighted, with red indicating peaks more accessible in cell lines and blue indicating peaks more accessible in liver.

**Figure 4.**
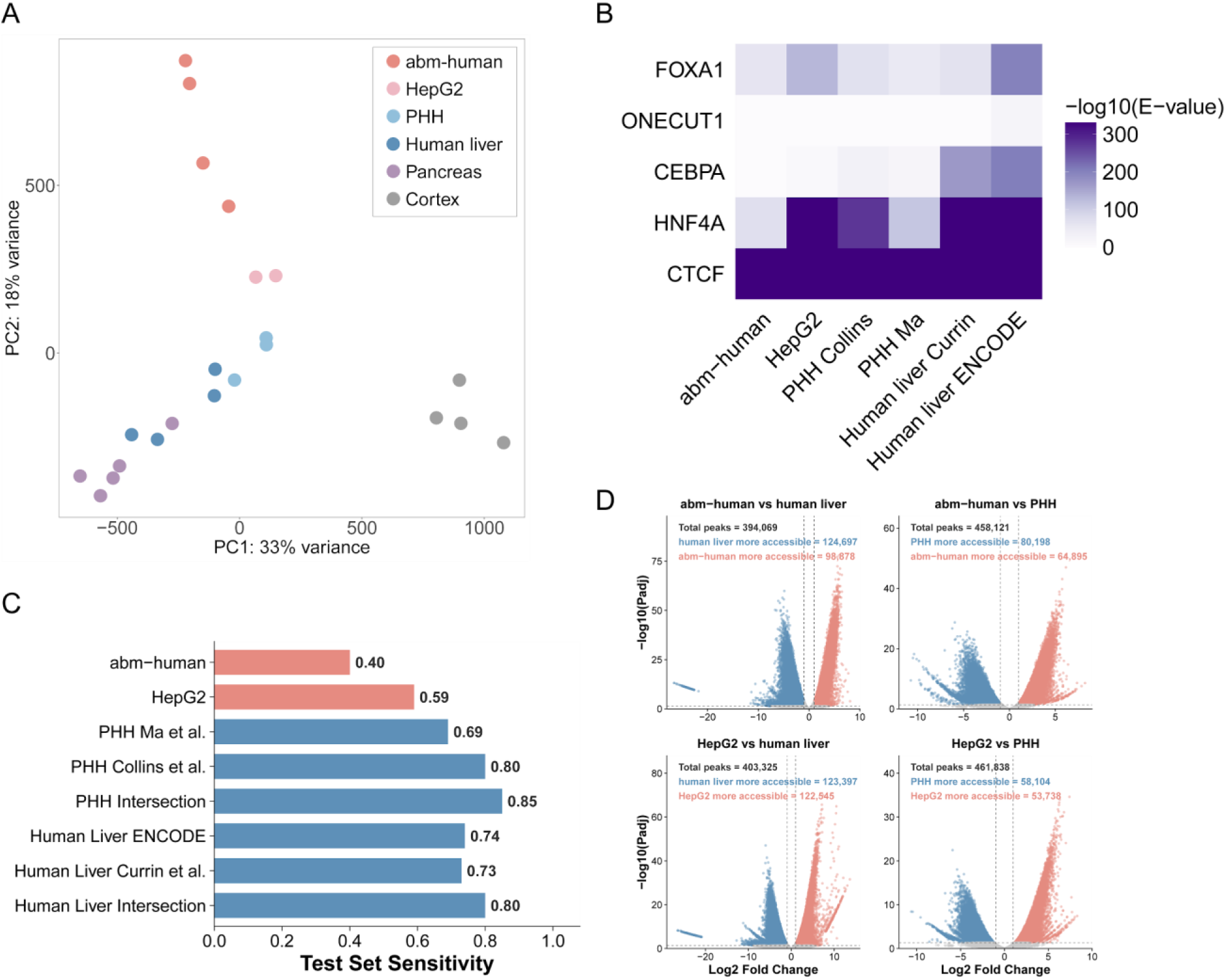
Liver-derived human cell lines show limited hepatocyte-like regulatory identity. **A)** Principal component analysis (PCA) of ATAC-seq from abm-human, HepG2, primary human hepatocytes (PHH), and bulk human liver, pancreas, and cortex. **B)** MEME-ChIP motif enrichment heatmap of enhancers from human liver-derived cell lines, PHH, and human liver datasets. Color indicates motif enrichment significance as -log10(E-value), where motifs that were not detected were assigned an E-value of 1. **C)** Sensitivity of liver machine learning model on human enhancers. **D)** DESeq2 differential accessibility volcano plots comparing ATAC-seq peaks between liver-derived human cell lines and reference hepatocyte/liver samples. Each point represents an accessible chromatin peak from comparisons of a human HLC (abm-human or HepG2) against bulk human liver or PHH. The x-axis shows log2 fold change in accessibility, and the y-axis shows -log10 adjusted p-value. Peaks with adjusted p < 0.05 and |log2 fold change| > 1 are highlighted, with red indicating peaks more accessible in cell lines and blue indicating peaks more accessible in liver or PHH.

Differential chromatin accessibility analysis also supported this divergence (**Figure 3C**). Enhancers more open in rat liver than in abm-rat were enriched for HNF4A, ONECUT1, and CEBPA motifs (**Tables S21-S22**). Genes near CREs more accessible in rat were enriched for occurring in biological processes related to hepatic metabolic functions, including xenobiotic metabolism and catabolism, monooxygenase activity, long-chain fatty acid metabolism, triglyceride metabolism, and P450-associated pathways. In contrast, genes near CREs more accessible in abm-rat were enriched for GO terms for non-hepatocyte programs, including apoptotic processes involved in morphogenesis, osteoclast and bone cell development, regulation of developmental apoptosis, collagen-activated signaling, and ribosomal and cytoskeletal categories (**Table S23**). Thus, like AML12, abm-rat cells seem to have retained only limited hepatocyte-like transcriptional regulatory features and lack the functional regulatory program seen in native rat liver.

We observed similar patterns in the human HLCs. We compared both abm-human and HepG2 cells with published ATAC-seq datasets from human liver tissue and primary human hepatocytes (PHH). PCA separated both cell lines from liver tissue and PHH, indicating substantial divergence from native hepatocyte chromatin accessibility landscapes, with abm-human showing greater divergence (**Figure 4A**). Liver tissue and PHH enhancers showed strong enrichment for HNF4A (human liver E-value = 5.7e-432 and 3.4e-340, PHH E-value = 5.40E-112 and 2.60E-276), CEBPA (human liver E-value = 2.1e-93 and 2.60E-169, PHH E-value = 1.80E-20 and 1.30E-31), and FOXA1 (human liver E-value = 7.7e-74 and 2.50E-66, PHH E-value = 1.50E-51 and 8.70E-69) motifs, whereas enrichment of these motifs was substantially reduced in both HLCs and most strongly depleted in abm-human (HepG2 HNF4A E-value = 9.3e-684, CEBPA E-value = 2.80E-9, FOXA1 E-value = 1.80E-131; abm-human HNF4A E-value = 1.90E-71, CEBPA not detected, FOXA1 E-value = 1.60E-60) (**Figure 4B**, **Tables S24-S29**). At the functional level, genes near liver and PHH CREs were enriched for GO terms related to hepatocyte-associated metabolic and endocrine-response programs, including gluconeogenesis, hexose or monosaccharide biosynthesis, glucose metabolism, cholesterol efflux or reverse cholesterol transport, insulin-response pathways, and thyroid hormone response. In contrast, genes near abm-human CREs were enriched for GO terms related to podosome, muscle alpha-actinin binding, TGF-β2 production, pericyte differentiation, and Notch target transcription. Genes near HepG2 CREs retained some enrichment for liver-related metabolic categories, but these were accompanied by non-canonical or less hepatocyte-specific terms including intestinal epithelial maturation, DNA integration, adherens junction maintenance, copper ion transport, and cytochrome-c oxidase activity (**Table S30**). Furthermore, the machine learning model for predicting liver enhancer activity achieved poor sensitivity on abm-human enhancers and slightly better than random sensitivity on HepG2 enhancers despite strong sensitivity for both primary human liver tissue enhancers and PHH enhancers (**Figure 4C**).

Differential accessibility analysis comparing each human HLC and primary liver tissue as well as PHH also showed widespread divergence. Approximately 60% of ATAC-seq peaks were differentially accessible between each HLC and bulk liver tissue, whereas about 30% differed relative to PHH, a result concordant with the cells having been derived specifically from hepatocytes (**Figure 4D**). Enhancers more accessible in liver tissue or PHH showed strong enrichment for HNF4A, CEBPA, and FOXA1 motifs. In contrast, enhancers more accessible in HLCs showed reduced or absent hepatocyte-associated motif enrichment, particularly for CEBPA, consistent with incomplete hepatocyte regulatory identity (**Tables S31-S38**). Genes near CREs more accessible in liver or PHH were enriched for GO terms related to core hepatic metabolic functions, including glucose, lipid, sterol, cholesterol, fatty acid, triglyceride, and alcohol metabolism. In contrast, genes near CREs more accessible in human HLCs were enriched for GO terms related to non-hepatocyte developmental, structural, migration-associated, and ectopic signaling programs (**Table S39**). Together, these results show that the human HLCs examined here also do not fully recapitulate the global chromatin accessibility patterns, enhancer motif landscapes, or functional transcriptional regulatory programs characteristic of native liver and primary hepatocytes.

## Discussion

Direct comparisons of CRE activity in immortalized cell lines to their tissues of origin have been limited despite the wide use of immortalized cell lines to test hypotheses regarding CRE activity(9–12). Previous studies have reported extensive gene expression differences between cell lines and their primary cell type of origin, but most focused on the comparison of cancer-derived cell lines(20) or blood and skin cell lines(16). Whether HLCs, which are widely used in CRE activity studies, preserve the chromatin accessibility landscapes needed to study liver CRE function has not previously been investigated. We therefore directly compared CRE activity as measured by ATAC-seq(56) between primary liver tissue and four HLCs including three non-cancer-derived cell lines: AML12, abm-rat, abm-human, and HepG2. Although each HLC’s CREs retained enrichment for some liver-associated TF motifs, all four HLCs showed reduced enrichment for at least one key liver TF motif in its CREs relative to primary liver tissue. In addition, CREs with stronger chromatin accessibility in primary liver tissue relative to HLCs were enriched for occurring near genes involved in core liver metabolic processes, while CREs with stronger chromatin accessibility in HLCs relative to primary liver tissue were enriched for occurring near genes involved in non-hepatic or less liver-specific processes, such as immune regulation, differentiation, and the cell cycle. Furthermore, a machine learning model trained to predict enhancer activity in primary liver tissue that yielded 82% sensitivity on parts of the genome not used in training and strong performance for its ability to predict enhancer activity differences between species(79) yielded less than 60% sensitivity on HLC enhancers. Together, these results indicate that HLCs do not fully recapitulate the transcriptional regulatory landscape of primary liver tissue. These results are concordant with a previous study showing extensive gene expression differences between PHH and three cancer-derived HLCs, including HepG2(20).

Interestingly, HepG2 was the HLC most similar to primary hepatocytes despite it being the only cancer-derived cell line included in our study. HepG2 was the only cell line with strong enrichment in CREs for the motif of HNF4A, a TF that plays essential roles in post-natal liver function(97,103–105). It was also the only HLC for which a machine learning model that accurately predicts primary liver tissue enhancer activity(79) achieved over 50% sensitivity.

These results are concordant with a previous study showing that HepG2 had higher expression of important liver gluconeogenesis genes G6PC and PCK1 than AML12 and THLE-2(51).

Together, these results suggest that HepG2 cells, despite their cancerous origins, may retain more liver regulatory and metabolic features than the other HLCs that we tested and thus be a more promising option for testing hypotheses regarding liver CRE activity.

The differences between human liver and HLCs are unlikely to be explained by the cellular heterogeneity of liver tissue or the culture conditions used for the cell lines. The proportion of significantly differentially accessible peaks was lower when comparing the human HLCs to PHH rather than bulk liver tissue, suggesting that differences between HLCs and PHH were not caused by HLCs coming from non-hepatocyte liver cell types. Although antibiotics have been shown to alter gene expression and CRE activity in HepG2(49,50), culturing each cell line without antibiotics for three weeks did not restore a liver-like chromatin accessibility landscape.

Likewise, growing AML12 cells without insulin for 36 hours did not increase their similarity to primary liver tissue. One possibility is that hepatocytes eventually lose mature hepatic identity when maintained outside the liver microenvironment, where they lack normal tissue architecture, extracellular matrix, and interactions with non-hepatocyte liver cells; this loss of cell identity would be concordant with previously reported difficulties in maintaining PHH in conventional cell culture(106–108).

The mechanisms underlying differences in CRE activity between HLCs and primary liver tissue are difficult to discern from this data alone and may differ between cell lines and culture conditions. One possibility is that immortalization ultimately causes a reversion of cells to a less mature state, which is supported by our finding that most AML12 cells had strong protein expression of immature hepatocyte marker CD24 but lacked mature hepatocyte marker ASGR1. Another possibility is that immortalization promotes proliferative regulatory programs. This is supported by our finding that CREs in multiple HLCs were enriched for occurring near genes involved in differentiation and previous investigations of blood and skin immortalized cell lines that obtained similar results(16). An additional possibility is that key liver TFs or their co-factors are insufficiently expressed in HLCs. While AML12 cells showed high expression of Hnf4a and Onecut1 in our RT-qPCR data, both TFs are known to interact with other TFs expressed in the liver(109–112) that could stabilize their binding. Thus, reduced expression or activity of these co-factors could weaken cooperative binding, limiting the retention of the mature liver regulatory landscape. RNA-seq in HLCs could help decipher such mechanisms. However, most publicly available HLC datasets were generated from cells grown in antibiotics(52) or under culture conditions that substantially differ from those recommended in the original description of the cell lines(31,97), so such experiments may need to be repeated under literature-supported optimal culture conditions.

Although we found substantial differences between HLCs and primary liver tissue, these do not mean that HLCs lack utility. First, we did not profile all available HLCs. However, previous studies and other available information suggest that the HLCs we did not test may also have limitations that affect their fidelity as models of primary liver tissue. For example, the abm HLCs not included in this study were immortalized using the same method as abm-rat(100), THLE-2 was previously shown to have a low protein abundance of the insulin receptor(51), and a previous study showed that multiple other cancer-derived HLCs differ from primary hepatocytes at the gene expression level(20). Second, we did not comprehensively explore cell culture conditions. Although the optimized culture method we used was based on previously published studies, a recent study reported that a modified AML12 cell culture protocol could improve its metabolic function, although publicly available experimental evidence is currently lacking(113). Thus, further optimization of cell culture conditions has the potential to increase the similarity between HLC and primary liver tissue CREs. Finally, even if HLCs have many differences in CRE activity from primary liver tissue, many CREs do overlap, and the subset of TF motif and GO enrichments that are shared may reflect regulatory programs that are partially retained in HLCs. In addition, they may still be useful for studying other components of the liver, such as insulin signaling, response to liver injury, or the effects of liver drugs, especially when the CREs or pathways under investigation are shown to be active in both the cell line and the relevant primary liver tissue context.

## Limitations of the study

We did most of our comparisons between cell lines and primary tissue using ATAC-seq data. Although ATAC-seq peaks are a proxy for CREs, chromatin accessibility is not an exact measure of CRE activity. For example, ATAC-seq can miss CREs due to biases in the assay and insufficient sequencing depth, and not all accessible chromatin regions act as CREs. In addition, we linked CREs to genes using the method described in GREAT, which creates regulatory domains for genes that are up to 1Mb in each direction from genes’ transcription start sites and links them to the CREs within them(95). Since this approach can cause incorrect CRE-gene links(114), our understanding of differences between up-regulated genes in cell lines and primary tissues may be inaccurate. This understanding may be even more limited because changes in activity of CREs do not always lead to changes in expression of the genes they regulate, as genes are often regulated by multiple CREs that could have different activity changes(115). A lack of change in CRE chromatin accessibility does not always indicate a lack of changes in gene expression, not only because genes can be regulated by multiple CREs but also because the binding of TFs in accessible chromatin regions can change without the openness of the region changing(116). Furthermore, changes in gene expression are far from perfectly indicative of changes in protein expression due to regulation at the translational level. Thus, comparisons of RNA-seq and MassSpec between multiple liver cell lines and primary liver tissue and PHH would be valuable extensions to this work. Finally, HLCs we did not test may have more similar transcriptional regulatory landscapes to primary hepatocytes than those we tested.

## Conclusion

Our findings of substantial differences between HLC and primary liver tissue CREs demonstrate the importance of comparing cell lines against primary tissues before using cell lines to test hypotheses about primary tissues. Ideally, such comparisons should be performed under optimized culture conditions and with assays that directly evaluate whether the cell line is suitable for the intended biological question. While cell lines can be valuable model systems because they reduce the cost, logistical challenges, and ethical concerns associated with animal models or primary tissue acquisition, assumptions should not be made about their fidelity to their tissue of origin. Instead, they should be rigorously characterized to assess their limitations and suitability for testing a given hypothesis. When such investigations suggest that cell lines are not useful proxies for primary tissue, *in vivo* assays, such as recently developed *in vivo* MPRAs(117–120) and CRISPR assays(121,122) , can be used as alternatives. These assays have been successful in multiple mouse tissues, including the liver(117,121). We therefore recommend that future investigations of CRE activity be carried out using a combination of different cell lines or *in vivo* experiments, where the combination should depend on the questions being asked.

## Description of Supplementary Tables

Table S1: RNA Integrity Numbers for RNA extracted for RT-qPCR experiments. All RINs were > 8.0, indicating highly intact RNA.

Table S2: MEME-ChIP results for AML12 cells cultured in complete medium Table S3: MEME-ChIP results for mouse liver Liu

Table S4: MEME-ChIP results for mouse liver Kaplow A

Table S5: rGREAT results for AML12 cells cultured in complete, -P/S, -insulin, -P/S -insulin medium

Table S6: MEME-ChIP results for regions with increased chromatin accessibility in AML12 cells cultured in complete medium compared with mouse liver

Table S7: MEME-ChIP results for regions with decreased chromatin accessibility in AML12 cells cultured in complete medium compared with mouse liver

Table S8: rGREAT results of differentially accessible chromatin regions identified by DESeq2 in AML12 cells cultured in complete, -P/S, -insulin, -P/S -insulin medium compared with mouse liver

Table S9: MEME-ChIP results for AML12 cells cultured in -P/S medium

Table S10: MEME-ChIP results for AML12 cells cultured in -P/S -insulin medium Table S11: MEME-ChIP results for AML12 cells cultured in -insulin medium

Table S12: MEME-ChIP results for regions with increased chromatin accessibility in AML12 cells cultured in -P/S medium compared with mouse liver

Table S13: MEME-ChIP results for regions with decreased chromatin accessibility in AML12 cells cultured in -P/S medium compared with mouse liver

Table S14: Table S12: MEME-ChIP results for regions with increased chromatin accessibility in AML12 cells cultured in -insulin medium compared with mouse liver

Table S15: Table S12: MEME-ChIP results for regions with decreased chromatin accessibility in AML12 cells cultured in -insulin medium compared with mouse liver

Table S16: MEME-ChIP results for regions with decreased chromatin accessibility in AML12 cells cultured in -P/S -insulin medium compared with mouse liver

Table S17: MEME-ChIP results for regions with increased chromatin accessibility in AML12 cells cultured in -P/S -insulin medium compared with mouse liver

Table S18: MEME-ChIP results for rat liver Stephen Table S19: MEME-ChIP results for abm-rat

Table S20: MEME-ChIP results for rat liver Takeuchi

Table S21: MEME-ChIP results for regions with increased chromatin accessibility in abm-rat cells compared with rat liver

Table S22: MEME-ChIP results for regions with decreased chromatin accessibility in abm-rat cells compared with rat liver

Table S23: rGREAT results of differentially accessible chromatin regions identified by DESeq2 in abm-rat cells compared with rat liver

Table S24: MEME-ChIP results for human liver ENCODE Table S25: MEME-ChIP results for human liver Currin Table S26: MEME-ChIP results for PHH Ma

Table S27: MEME-ChIP results for PHH Collins Table S28: MEME-ChIP results for HepG2

Table S29: MEME-ChIP results for abm-human

Table S30: rGREAT results of HepG2, abm-human, human liver ENCODE, human liver Currin, PHH Ma, PHH Collins

Table S31: MEME-ChIP results for regions with decreased chromatin accessibility in HepG2 compared with human PHH

Table S32: MEME-ChIP results for regions with decreased chromatin accessibility in abm-human compared with human PHH

Table S33: MEME-ChIP results for regions with increased chromatin accessibility in HepG2 compared with human PHH

Table S34: MEME-ChIP results for regions with increased chromatin accessibility in abm-human compared with human PHH

Table S35: MEME-ChIP results for regions with decreased chromatin accessibility in HepG2 compared with human liver

Table S36: MEME-ChIP results for regions with increased chromatin accessibility in HepG2 compared with human liver

Table S37: MEME-ChIP results for regions with increased chromatin accessibility in abm-human compared with human liver

Table S38: MEME-ChIP results for regions with decreased chromatin accessibility in abm-human compared with human liver

Table S39: rGREAT results of differentially accessible chromatin regions identified by DESeq2 in HepG2 and abm-human compared with human liver and PHH

## List of Abbreviations

CRE: Cis-regulatory element
HLC: hepatocyte-like cells
AML12: Alpha mouse liver 12 cell line
MPRA: massively parallel reporter assay
TF: Transcription factor
RNA-seq: RNA Sequencing
THLE-2: Telomerase-immortalized Liver Epithelial-2 cell line
ATAC-seq: Assay for Transposase-Accessible Chromatin with sequencing
abm-rat: Immortalized rat hepatocyte cell line from Applied Biological Materials, Inc, Catalog # T0078
abm-human: Immortalized human hepatocyte cell line from Applied Biological Materials, Inc, Catalog # T0063
GO: Gene Ontology
P/S: penicillin/streptomycin
PCA: principal component analysis
PHH: primary human hepatocytes
DMEM: Dulbecco’s Modified Eagle Medium
ITS: Insulin/transferrin/selenium
RT-qPCR: Reverse Transcription-Quantitative Polymerase Chain Reaction

## Declarations

### Ethics approval and consent to participate

All human data is from post-mortem samples and was made publicly available by previous studies that were conducted in accordance with NIH guidelines. Thus, none of the accompanying data included in this manuscript comes from living subjects, nor is there de-identifiable information on any of the subjects. All data from non-human animals is either publicly available or was collected in experiments approved by Carnegie Mellon University’s Institutional Animal Care and Use Committee and reported in accordance with the ARRIVE guidelines.

### Consent for Publication

Not applicable.

### Availability of data and materials

Data presented in this study is available through the National Center for Biotechnology Information Gene Expression Omnibus (GSE334599), and the reviewer token is yjijuykktfkbvcf. Publicly available data used in this study can be found as described in the **Methods** section.

Code created for this study can be found at https://github.com/KaplowLab/Comparison_of_liver_cell_line_and_native_liver (Zenodo: 10.5281/zenodo.20544805). All other code and the machine learning model used in this study can be found as described in the **Methods** section.

### Competing interests

The authors declare no competing interests.

### Funding

This research did not receive any specific grant from funding agencies in the public, commercial, or not-for-profit sectors.

## Acknowledgements

We would like to thank A. R. Brown for providing mouse liver tissue for RT-qPCR and K. Mohlke for providing information about the human liver samples from (60). We would also like to thank E. Guilfoyle, C. Key, and the other members of the Kaplow Lab for useful discussions and suggestions.

## Authors’ Contributions

Conceptualization: A.B. and I.M.K.; methodology: A.B., I.M.K.; software: X.L., D.M.-F., D.X., G.V., J.M., I.M.K.; validation: A.B., X.L., D.M.-F., G.V., Y.C., I.M.K.; formal analysis: A.B., X.L., D.M-F., G.V., J.M., I.M.K.; investigation: A.B., X.L., D. M.-F., D.X., K.D., G.V., J.M., B.A.A.E., M.K., Y.C.; resources: A.B., J.M., I.M.K.; data curation: X.L., D. M.-F., G.V., J.M., I.M.K.; writing original draft: A.B., X.L., D. M.-F., I.M.K.; writing review and editing: all authors; visualization: A.B., X.L., D.M.-F., J.M., A.O.; supervision: A.B., J.M., I.M.K.; project administration: A.B., I.M.K.

## Corresponding author

Requests for further information and resources should be directed to and will be fulfilled by the lead contact, Irene M. Kaplow.

## Notes

https://github.com/KaplowLab/Comparison_of_liver_cell_line_and_native_liver

### Competing Interest Statement

The authors have declared no competing interest.

### Summary of Updates

We revised the Abstract and added more details to the Methods.

