## Supplementary material for "Deciphering the limitations of immortalized hepatocyte cell lines for the study of liver cis-regulatory elements": Fig S1

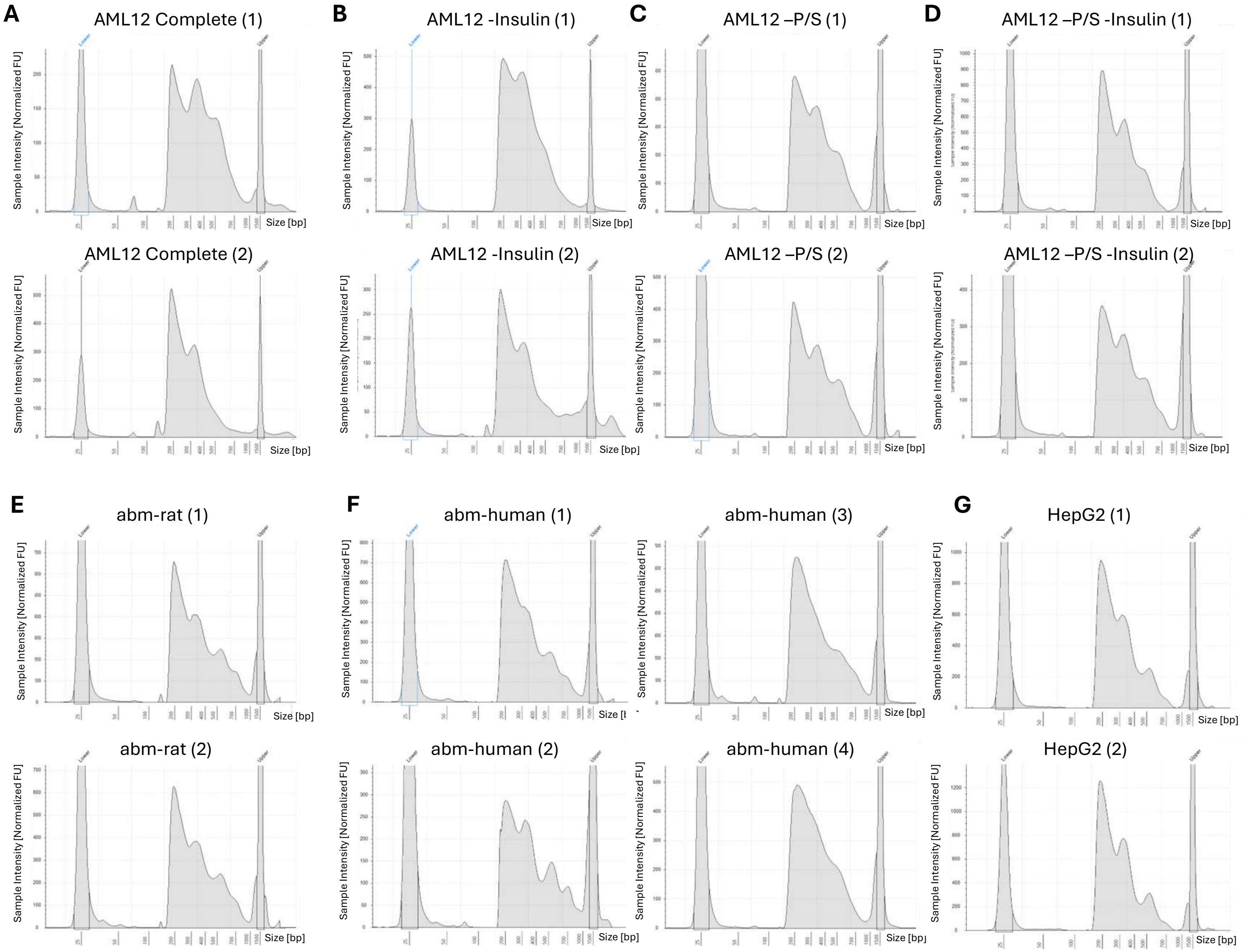
A
AML12 Complete (1)
AML12 Complete (2)
B
AML12 -Insulin (1)
AML12 -Insulin (2)
C
AML12 –P/S (1)
AML12 –P/S (2)
D
AML12 –P/S -Insulin (1)
AML12 –P/S -Insulin (2)
E
abm-rat (1)
abm-rat (2)
F
abm-human (1)
abm-human (2)
abm-human (3)
abm-human (4)
G
HepG2 (1)
HepG2 (2)
Sample Intensity [Normalized FU]
Size [bp]
